# Overexpression of NDR1 Leads to Pathogen Resistance at Elevated Temperatures

**DOI:** 10.1101/2021.12.01.470751

**Authors:** Saroopa P. Samaradivakara, Huan Chen, Yi-Ju Lu, Pai Li, Yongsig Kim, Kenichi Tsuda, Akira Mine, Brad Day

## Abstract

Abiotic and biotic environments influence a myriad of plant-related processes, including growth, development, and the establishment and maintenance of interaction(s) with microbes. As a driver of this signaling between plants and microbes, the role of plant hormones in both surveillance and signaling has emerged as a point of intersection between plant-abiotic and -biotic responses. In the current study, we elucidate a role for NON-RACE-SPECIFIC DISEASE RESISTANCE1 (NDR1) by exploiting effector-triggered immunity (ETI) to define the regulation of plant host immunity in response to both pathogen infection and elevated temperature. We generated time-series RNA sequencing data of WT Col-0, a NDR1 overexpression line, as well as ndr1 and ics1-2 mutant plants under elevated temperature. Not surprisingly, the NDR1-overexpression line showed genotype-specific gene expression changes related to defense response and immune system function. Interestingly, overexpression of NDR1 revealed a role for NDR1 in immune system function; specifically, we describe a mechanism that intersects with *Pseudomonas syringae*, type-III effector translocation, R-protein signaling complex stabilization, and sustained levels of SA at elevated temperature during ETI. The results described herein support a role for NDR1 in maintaining cell signaling during simultaneous exposure to elevated temperature and avirulent pathogen stressors.

**One-sentence summary:** NDR1 is required for *Pst*-AvrRpt2 triggered ETI at elevated temperature.

## INTRODUCTION

Plant response to abiotic and biotic stress requires the coordinated activity of numerous cellular processes, the vast majority of which share overlapping functions in basic physiological programs, including growth, development, and reproduction (Nejat and Mantri, 2017; Saijo and Loo, 2020). In recent years, the impact of elevated temperature on plant growth and defense has received increasing attention, due in part to ongoing changes in global climate and environmental stress (Havko et al., 2020; Zhao et al., 2020; Zhang et al., 2021). However, the precise mechanisms that govern immunity at elevated temperature remain undefined.

In large part, plant growth, development, and immune signaling processes are each influenced by fluctuations in temperature and environment (Zhu et al., 2010; Cheng et al., 2013; Bahuguna and Jagadish, 2015), the outcome of which is a reduction in vegetative plant growth (Quint et al., 2016), impacts on flower development and fertility (Balasubramanian et al., 2006; Koini et al., 2009; McClung and Davis, 2010), and the inhibition of plant defense signaling in response to a range of biotic threats (Wang et al., 2009; Wang and Hua, 2009). Not surprisingly, plants have evolved mechanisms to cope with simultaneous exposure to biotic and abiotic stress, and in this, utilize overlapping mechanisms to not only respond to stress, but to anticipate environmental changes for the purpose of regulating the timing and amplitude of seemingly opposing signaling processes (Quint et al., 2016; Gimenez et al., 2018; Saijo and Loo, 2020; Iqbal et al., 2021).

As a point of convergence with biotic stress signaling, changes in the abiotic environment have been shown to profoundly impact the plant immune system, including the activation, duration, and attenuation of signaling (Venkatesh and Kang, 2019). Indeed, recent studies have demonstrated that the function of at least two key nodes of the plant immune system – namely, pathogen associated molecular pattern (PAMP)-triggered immunity (PTI) and effector-triggered immunity (ETI) – are intimately associated with processes required for response to abiotic stress (Tsuda et al., 2009). ETI, which is manifested following the recognition of pathogen race-specific avirulence (Avr) proteins (aka, effectors), is regulated by host plant-derived resistance (R) genes (Jones and Dangl, 2006). As a highly conserved family of proteins found in all plants, nucleotide binding-leucine rich repeat (NB-LRR) protein molecules mediate the specific recognition of pathogens via the indirect and/or direct recognition of both conserved and race-specific virulence factors (Elmore et al., 2011).

In addition to NB-LRR proteins, numerous additional proteins and processes have been identified as critical components of the immune signaling network (De Vleesschauwer et al., 2014; Tsuda and Somssich, 2015; Li and Day, 2019; Maier et al., 2021). Interestingly, research has also demonstrated a role for these (i.e., phytohormones, transcription factors) in abiotic stress signaling (Berens et al., 2019; Saijo and Loo, 2020). Among these, *NON-RACE-SPECIFIC DISEASE RESISTANCE-1* (*NDR1*) was identified nearly three decades ago as a critical component of plant immune system function (Century et al., 1995), with key functions associated with ETI- and salicylic acid (SA)-dependent, signaling networks in Arabidopsis (Lu, 2009; Lu et al., 2013). As a broader role for NDR1 in plant processes, recent work has shown that NDR1 and NDR1-like genes (i.e., HIN; (Bao et al., 2016) play important roles in stress response signaling (Lu et al., 2021). Among the best characterized examples of *NDR1*-dependent immune signaling cascades is *RESISTANCE TO PSEUDOMONAS SYRINGE-2 (RPS2)* (Kunkel et al., 1993), a NB-LRR-encoding gene required for the recognition and activation of resistance in response to the Gram-negative bacterial phytopathogen *P. syringae* expressing the type III effector (T3E) protein AvrRpt2. In the absence of RPS2, AvrRpt2 promotes pathogen virulence in host cells (Mudgett, 2005). As a function for the role of NDR1 in RPS2 signaling, previous work demonstrated that RPS2-mediated resistance is *NDR1*-dependent (Axtell et al., 2003).

Herein, we describe a role for *NDR1* in plant immunity under heat stress conditions. We have identified a genotype specific cluster of highly expressed genes in the *NDR1*-overexpression line under heat stress. Most notably, unlike the temperature sensitive SA defense pathway gene *ISOCHRISMATE SYNTHASE 1* (*ICS1*) (Huot et al., 2017), the *NDR1*-overexpression line stabilizes ETI-specific *RPS2* mRNA accumulation at elevated temperature. Our findings suggest pathogen resistance at elevated temperature is mediated through crosstalk between NDR1 and RPS2, a mechanism that requires robust signaling of SA processes.

## RESULTS

### Temporal Dynamics of Transcriptome Responses to Heat Stress Through NDR1-Dependent Immune Activation

The loss of *NDR1* has a profound impact on pathogen defense signaling and disease resistance in plants (Century et al., 1995). Previous results suggest that one mechanism underpinning this activity may intersect with broader stress response processes, including those associated with plant hormone-based signaling and the maintenance of cellular integrity (Knepper et al., 2011). To define how *NDR1* influences plant response to abiotic stress response, we first conducted a comprehensive RNA-seq analysis over a 24 h period following permissive temperature (i.e., 21ºC) and heat stress (i.e., 29ºC) exposure in WT Col-0, the *ndr1* mutant, a previously characterized *ndr1/35S::NDR1*-overexpression line (Coppinger et al., 2004), and the SA deficient mutant *ics1-2*. The impetus for this was to determine the rapid transcriptional responses required for signaling in response to pathogen infection and elevated temperature, as well as to define the potential priming of immune responses and their relationship to heat tolerance.

As shown in Figure 1A, hierarchical clustering analysis of 11,245 differentially expressed genes (DEGs) revealed gene expression changes over the 24 hour (h) time course across all genotypes (Supplemental Data Set 1). Further analysis identified 2 significant gene clusters that showed significant response(s) to heat stress. The first, Cluster 1, contains 800 genotype-independent temperature responsive genes (Figure1A, Supplemental Figure 1A, Supplemental Data Set 2 and 3). As revealed by gene ontology (GO) enrichment analysis, this cluster contains a large number of genes involved in mitochondrial RNA editing, suggesting the role of mitochondrial RNA editing in acclimation to high temperature. This is supported by a recent paper reporting that an Arabidopsis mutant lacking the mitochondrial RNA editing enzyme, GEND1, is hypersensitive to high temperature (Guo et al., 2021). Notably, Cluster II, which is comprised of 2151 genes, is enriched in transcripts that were highly expressed in the *NDR1*-overexpression line (*ndr1/35S::NDR1*) and are related to defense response and immune system function based on GO enrichment analysis (Figure 1B, Supplemental Data Set 4 and 5). Interestingly, following 24 h exposure to elevated temperature, *NDR1*-overexpressing plants had the greatest number of DEGs, up- or downregulated, compared to *ndr1* and *ics1-2* (with WT Col-0 as a baseline) (Figure 1C, Supplemental Data Set 6 and 7). Taken together, overexpression of *NDR1* imparts a preemptive activation of immunity by heat stress.

**Figure 1.**
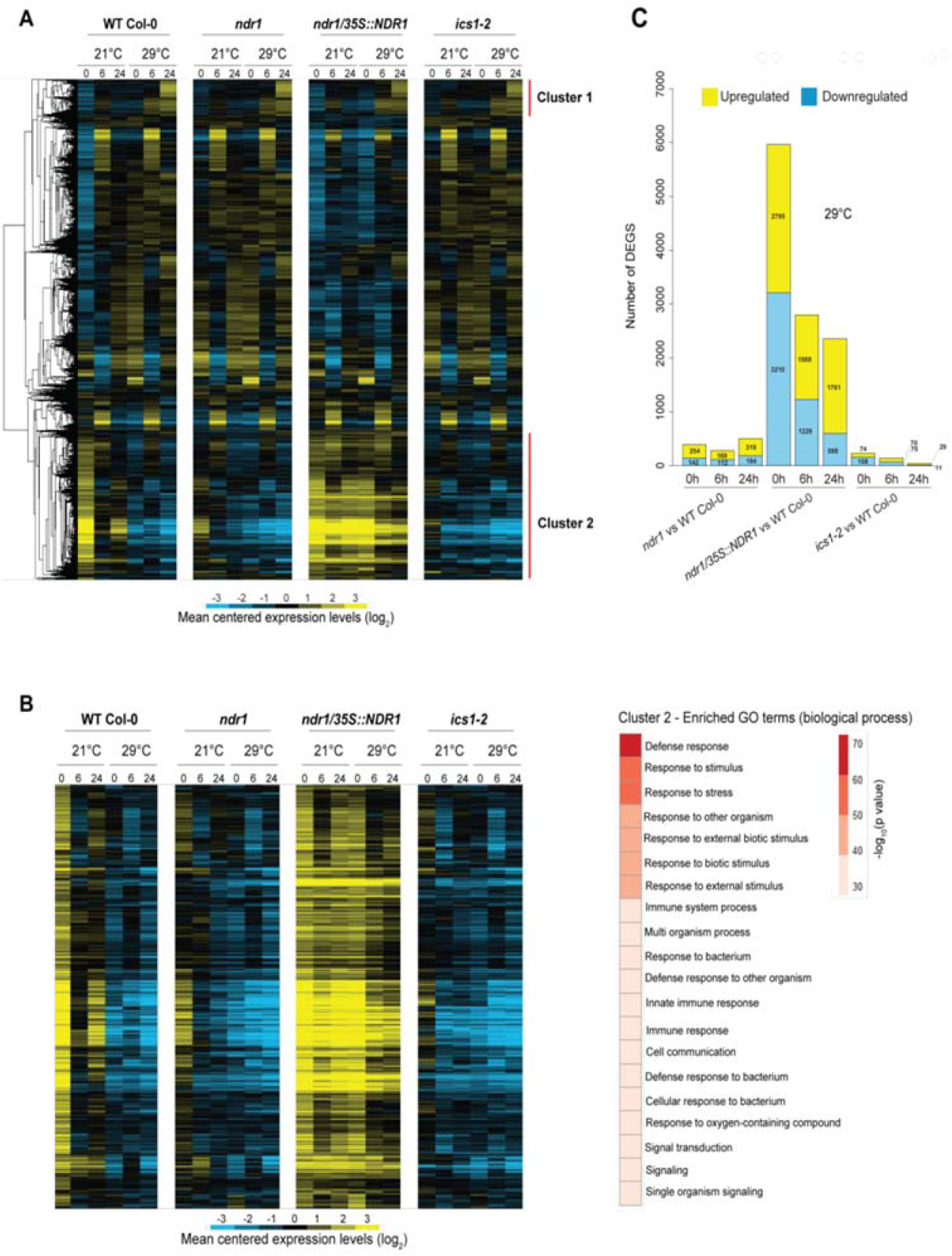
Temporal dynamics of transcriptome responses to heat stress through NDR1-dependent immune activation. **(A)** Heat map showing log_2_-fold gene expression changes over the 24 h heat stress. **(B)** Heat map showing log_2_-fold gene expression changes in Cluster II genes that are highly expressed in *ndr1/35S::NDR1* plants and is related to defense response and immune system based on GO enrichment analysis. Blue indicates negative values, yellow indicates positive values, and black indicates zero. **(C)** Number of differentially-expressed genes (DEGs) at 29°C over a 24 h time course in the *ndr1, NDR1*-overexpression and *ics1-2* mutant plants.

### Overexpression of *NDR1* Results in Sustained Accumulation of *RPS2* mRNA

The data described above support a role for transcriptional induction of defense responses in the *NDR1*-overexpression line at elevated temperature. This is exciting, as it points to a possible intersection between immunity and elevated temperature response through *NDR1*, a key regulator of ETI-based immune activation and signaling. To gain insight into the role of NDR1 at the intersection of immunity and high temperature response, we performed a co-expression network analysis using the R package WGCNA. This approach led to the identification of 37 modules with distinct expression patterns, as indicated by module eigengenes (MEs), which summarized the expression levels of the corresponding modules (Supplemental Figure 2). Using this, we calculated correlations of the expression pattern of *NDR1* and those of MEs. From this, we selected highly correlated modules (|correlation coefficient| > 0.6) to construct an *NDR1*-centered co-expression network (Figure 2A). Within this network, *NDR1* showed positive and negative correlations with modules 1 and 6 and modules 2 and 4, respectively. Modules 1 and 6 showed upregulation in the *NDR1*-overexpression line and this upregulation was maintained at elevated temperature (Figure 2B). Further, these modules were enriched for genes associated with immunity-related GO terms, such as “defense response” and “innate immune response” (Figure 2A and Supplemental Dataset 8 and 9). In contrast, modules 2 and 4 showed heat-resistant downregulation in the *NDR1*-overexpression line and were enriched for genes associated with photosynthesis- and growth-related GO terms (Figure 2A and Supplemental Dataset 8 and 9). The output of this analysis revealed that *NDR1* overexpression activates defense-associated gene expression and protects these expression networks from perturbation by elevated temperature.

**Figure 2.**
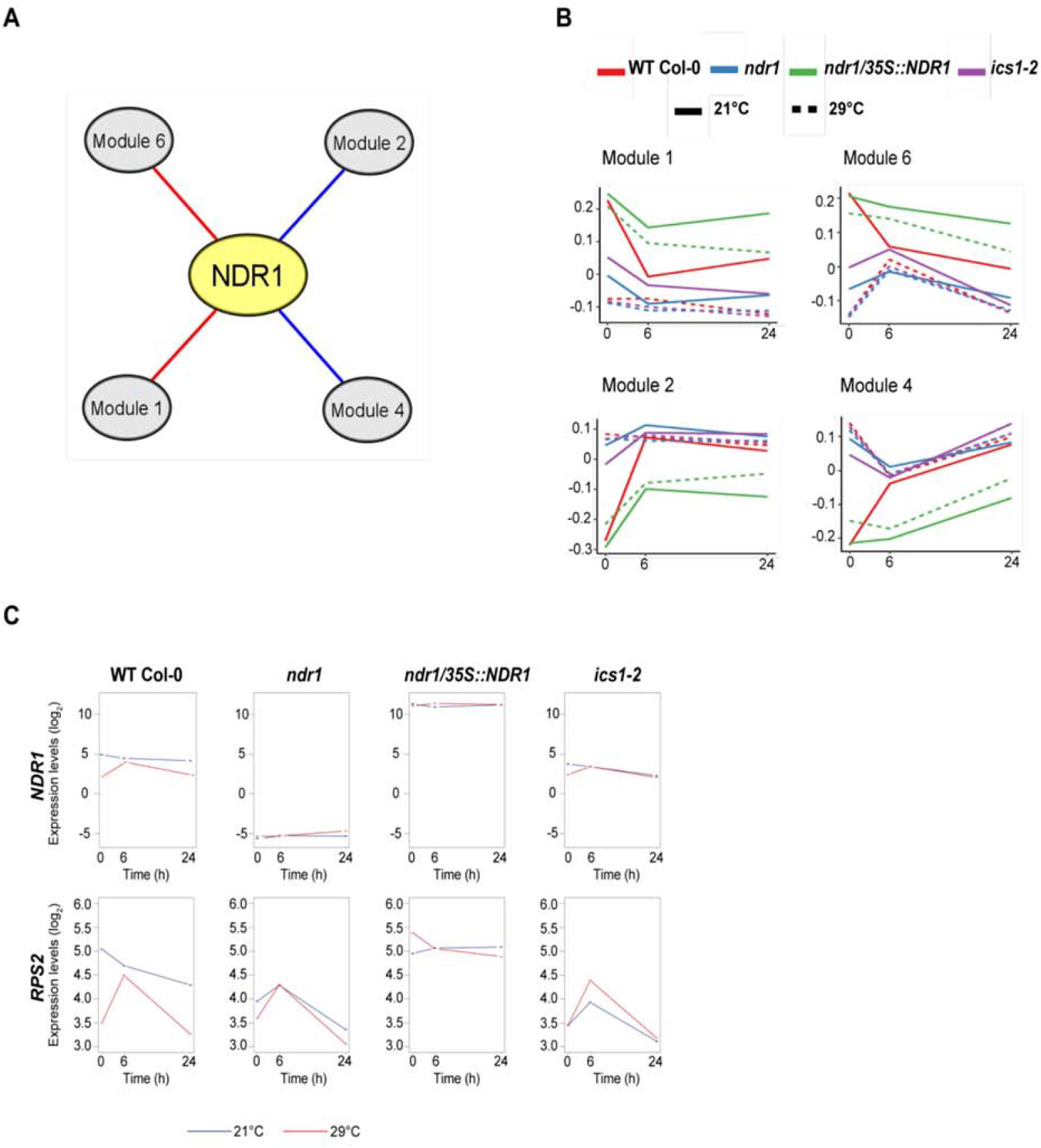
*NDR1* overexpression induces expression of immunity genes and protects it from perturbation by elevated temperature. **(A)** An *NDR1*-centered co-expression network reveals modules whose expression levels are correlated at elevated temperature in the *NDR1*-overexpression line. Red and blue edges indicate positive and negative correlation, respectively. **(B)** Averaged expression levels of genes in the modules summarized by module eigengenes. **(C)** Overexpression of *NDR1* results in sustained accumulation of *RPS2* mRNA at both 21°C and 29°C.

*NDR1* is required for the activation of ETI through a defined set of NB-LRR R-proteins (e.g., RPS2, RPM1) (Day et al., 2006; van Wersch et al., 2020). Previous studies showed that RPS2 is required for *Psm* ES4326 AvrRpt2- and *Pto* DC3000 AvrRpt2-induced SA accumulation and the induction of immune-associated transcripts (Liu et al., 2016; Mine et al., 2018). Interestingly, we found that *RPS2* is included in Module 6 (i.e., heat-resistant upregulation) in the *NDR1*-overexpression line. Based on this, we further evaluated the mRNA accumulation of *RPS2* and other key defense-associated genes at both permissive (21ºC) and elevated (29ºC) temperatures (Figure 2C and Supplemental Figure 1B). In contrast to the other genotypes, the downregulation of *RPS2* and *NDR1* (control) at 24 h was not observed in the *NDR1*-overexpression line at elevated temperature (Figure 2C and Supplemental Data Set1). Expression of the key genes in SA response, *ICS1, CALMODULIN BINDING PROTEIN 60g* (*CBP60g)*, and *PATHOGENESIS RELATED GENE 1* (*PR1*) was reduced at elevated temperature, but still higher in the NDR1-overexpression line as compared to the other genotypes (Supplemental Figure 1B). The gene expression level changes observed at T_0_ in all genotypes at 21ºC and 29ºC is likely due to occur through a combination of factors, such as the function of the genes themselves, changes occurring in response to the transfer of plants from permissive to elevated temperature chamber, and/or due to wounding during sampling.

To further define NDR1*’s* role as a regulator of general stress response signaling in Arabidopsis, we next asked if NDR1 is required for disease resistance signaling at elevated temperature. To do this, we first evaluated the activation of immune signaling in response to simultaneous exposure of elevated temperature and pathogen infection. Consistent with the requirement for NDR1 in the activation of RPS2-mediated ETI, WT Col-0 and *ndr1*/*35S*::*NDR1*, but not *ndr1*, responded to *Pst*-AvrRpt2 with rapid induction of the hypersensitive response (HR) at both permissive and elevated temperatures (Figure 3A, top 2 panels). This result was consistent with the absence of disease symptoms in WT Col-0 and the *NDR1*-overexpression line, and the development of disease symptoms (e.g., chlorosis) (Figure 3A, lower 2 panels) in both the *ndr1* and *ics1-2* mutant. As a further confirmation of this interaction, we also evaluated the *in planta* bacterial growth at 3-days after inoculation (DAI) to examine the level of host resistance and/or susceptibility against *Pst* DC3000 (*Pst*) and *Pst-*AvrRpt2. As shown, and consistent with the results of the HR assay, we observed enhanced susceptibility in plants lacking *NDR1* (*ndr1*) and SA (*ics1-2*), while WT Col-0 and *ndr1*/*35S*::*NDR1* showed resistance at elevated temperature (Figure 3B). *In planta* bacterial growth at 0 h post inoculation (hpi) was also quantified to capture any population-dependent growth rate differences (Supplemental Figure 3). Collectively, these results demonstrate that key regulators of SA, as well as the expression of *NDR1*-dependent resistance signaling (e.g., RPS2), are enhanced in the *NDR1*-overexpression line at elevated temperature.

**Figure 3.**
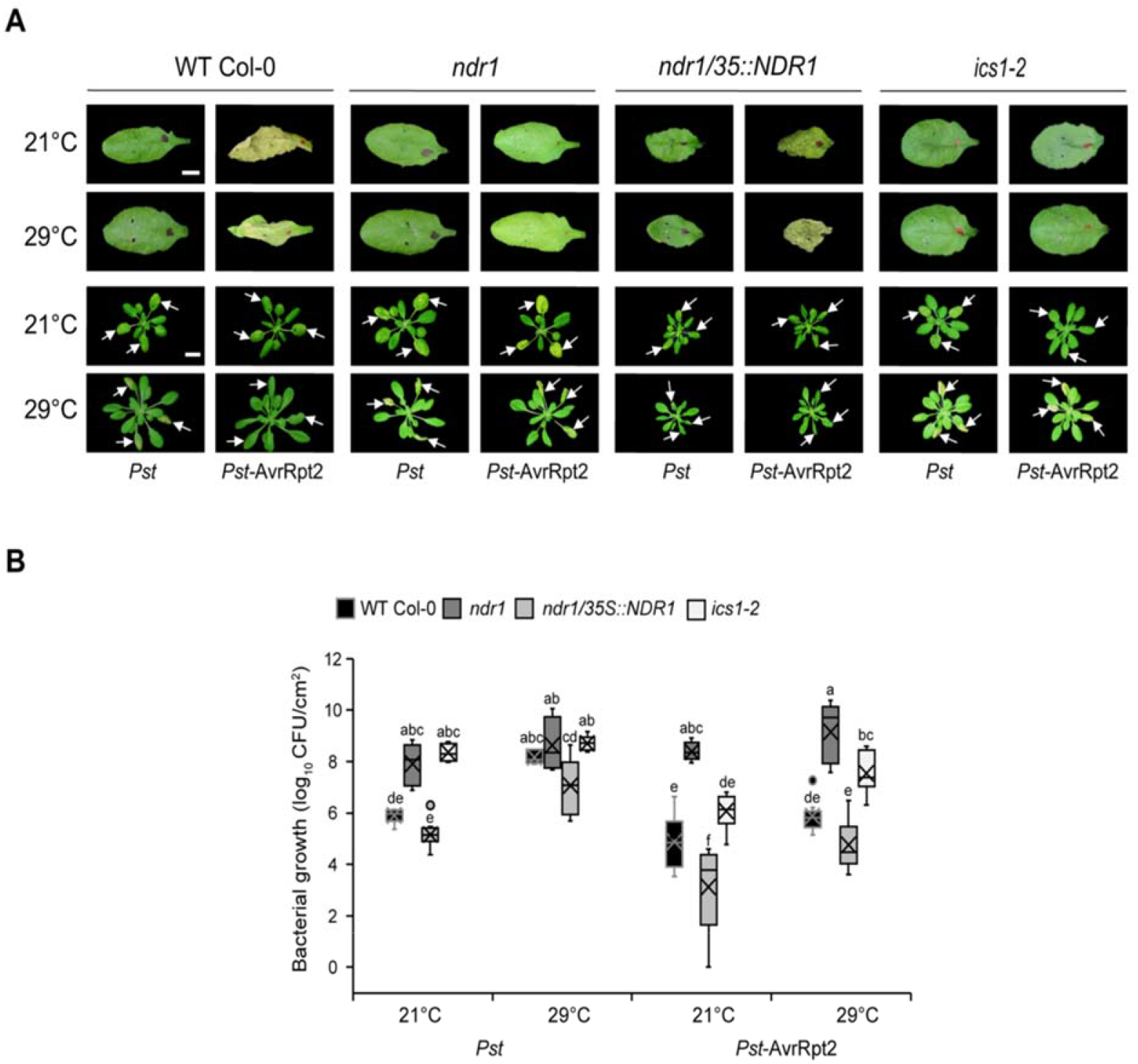
Disease resistance at elevated temperature is linked to stable *RPS2* mRNA expression in *ndr1/35S::NDR1* plants. **(A)** Hypersensitive response at 24 hpi (OD_600nm_ = 0.1) (top panel) and disease symptoms (OD_600nm_ = 0.0005) at three dpi (bottom panel) at 21°C and 29°C. Arrows indicate inoculated leaves. **(B)** Bacterial growth at 3 days after syringe-infiltration with *Pst* and *Pst*-AvrRpt2 (OD_600nm_ = 0.0005) in wild-type (Col-0) and mutant plants at 21°C and 29°C. *n* represents the total number of leaves from three independent biological repeats (n = 9). Values are plotted as box plots split by the median. Different letters represent a significant difference at *P* < 0.05 with Tukey’s honest significant difference (HSD) test. Bar = 0.5 cm. All data are representative of three independent experiments.

### *Pst*-AvrRpt2 promotes ETI-induced SA Accumulation at Elevated Temperatures

To determine if the observed disease resistance phenotype in the *NDR1*-overexpression line following challenge with avirulent *Pst*-AvrRpt2 is mediated by SA at elevated temperatures, we quantified the level of SA in plants hand-infiltrated with *Pst* and *Pst*-AvrRpt2 at 24 hpi. Consistent with previous reports, we observed a decreased SA accumulation at elevated temperature, compared to those at permissive temperature (21°C), following mock and *Pst* treatment (Figure 4A and B) (Huot et al., 2017). Intriguingly, we found that ETI triggered by *Pst*-AvrRpt2 led to a significant increase in the levels of SA in WT Col-0 (Figure 4B). In addition, the SA levels in the *ndr1/35S::NDR1*-overexpression line remained stable, in comparison to plants inoculated with the virulent pathogen *Pst* at elevated temperature (Figure 4C). The low levels of SA in the *ndr1* and *ics1-2* mutant plants are consistent with the observed susceptibility to *Pst*-AvrRpt2 at both permissive and elevated temperatures (Figure 3B).

**Figure 4.**
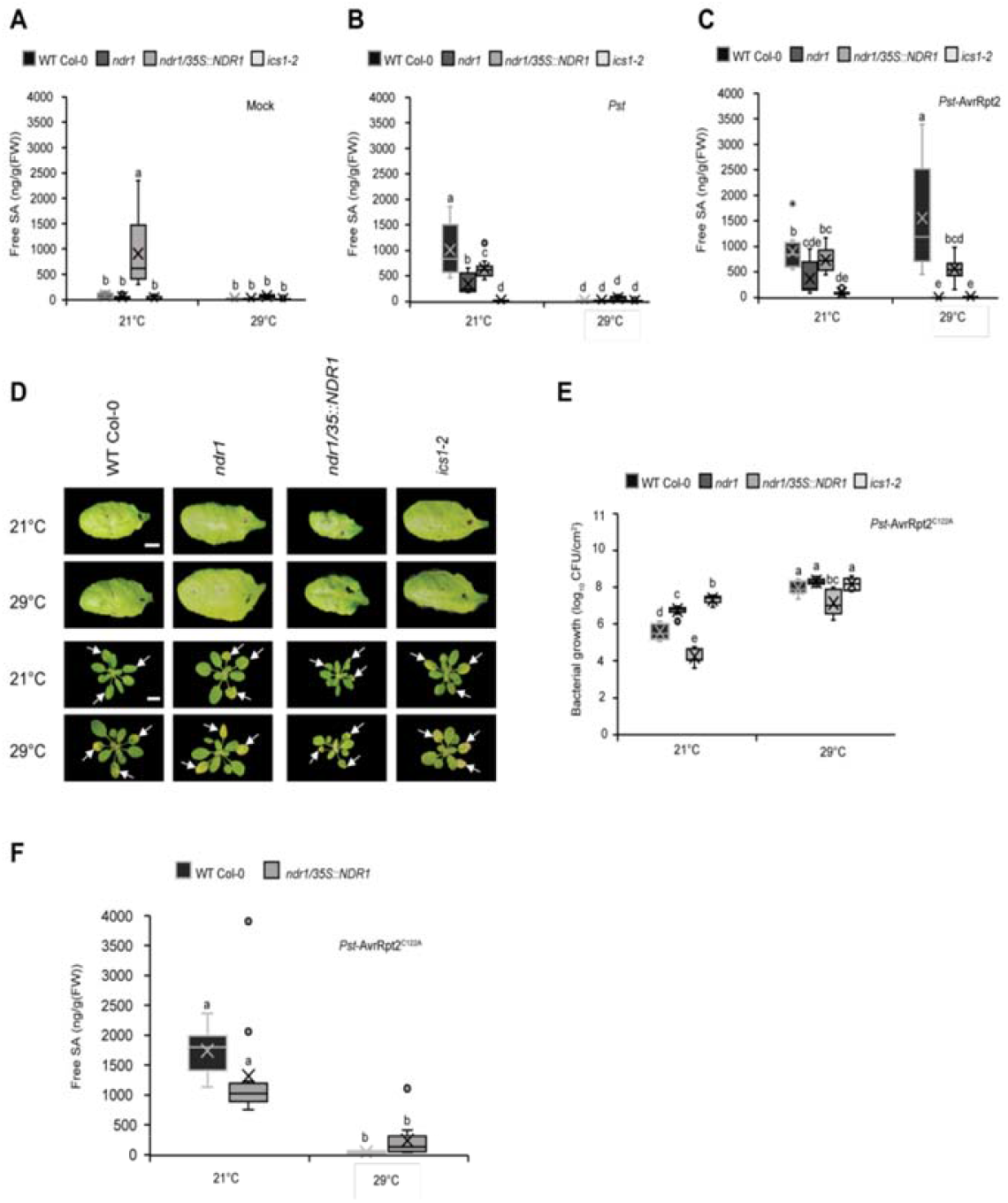
*Pst*-AvrRpt2 promotes ETI induced SA synthesis at elevated temperature. **(A)** Basal accumulation of total SA in wild-type (Col-0) and mutant plants. Leaves of mock infiltrated plants were harvested at 24 hpi and evaluated for SA content. **(B)** Pathogen-induced SA levels in wild-type (Col-0) and mutant plants treated with *Pst* and **(C)** *Pst*-AvrRpt2 (OD_600nm_ = 0.0005). Leaves of pathogen-infiltrated plants were harvested at 24 hpi for SA quantification. **(D)** Hypersensitive response (HR) at 24 hpi (OD_600nm_ = 0.1) (top panel) and disease symptoms (OD_600nm_ = 0.0005) at 3 dpi (lower panel) after syringe-infiltration with *Pst*-AvrRpt2^C122A^ in wild-type (Col-0) and mutant plants. **(E)** Bacterial growth at 3 days after syringe-infiltration with *Pst-AvrRpt2*^C122A^ (OD_600_=0.0005) in wild-type (Col-0) and mutant plants. **(F)** Levels of free SA in WT Col-0 *and ndr1/35S::NDR1* plants treated with *Pst*-AvrRpt2^C122A^ (OD_600_=0.0005). *n* represent total number of leaves from three independent biological repeats (For hormone quantification and disease assays n = 12 and 9, respectively). Measures are plotted as box plots split by the median. Different letters represent a significant difference at *P* < 0.05 with Tukey’s honest significant difference (HSD) test. Bar = 0.5 cm. All data are representative of three independent experiments.

To determine if overexpression of *NDR1*, and/or *Pst*-AvrRpt2 infection, is responsible for the induction of SA at elevated temperature, we first evaluated the *in planta* bacterial growth in plants infiltrated with *Pst*-AvrRpt2, and the AvrRpt2 cysteine protease mutant AvrRpt2^C122A^ (Kim et al., 2005). As expected, in the absence of the cysteine protease activity of AvrRpt2 (e.g., *Pst* alone or AvrRpt2^C122A^), we observed the absence of HR elicitation and the development of pronounced disease phenotypes in all plant lines at both permissive and elevated temperatures (Figure 4D and E). Next, to define the link between the cysteine protease activity of AvrRpt2 and the induced accumulation of SA, we quantified the level of SA in WT Col-0 and the *NDR1*-overexpression plants at 24 hpi with *Pst*-AvrRpt2^C122A^. In contrast to elevated levels of SA in plants following *Pst*-AvrRpt2 infection, we observed low levels of SA at elevated temperature in WT Col-0 and *NDR1*-overexpression plants following AvrRpt2^C122A^ inoculation (Figure 4F). Coupled with the results above (Figure 4C), these data support a role for *Pst*-AvrRpt2-associated temperature independent SA levels in WT Col-0 or the *NDR1*-overexpression plants grown at elevated temperatures. Based on this, we hypothesize that the observed resistance at elevated temperature is mediated by *Pst-*AvrRpt2 induced SA production/stabilization, as well as through *NDR1*-overexpression.

### Overexpression of *NDR1* Leads to Enhanced Stability of RIN4 in the Presence of *Pseudomonas syringae* Expressing *Pst-*AvrRpt2

The data presented above support a link between the immunity-associated function of NDR1 at elevated temperature and pathogen-induction of SA. To further define the mechanism(s) underpinning this activity, we next investigated the activation of ETI through the NDR1-RIN4 signaling node. We first evaluated the activity of the T3E cysteine protease AvrRpt2, by examining its ability to cleave RIN4 (Axtell et al., 2003; Chisholm et al., 2005). To begin, we quantified the RIN4 protein levels of the untreated plants at both permissive and elevated temperatures. The RIN4 protein levels were similar at both temperatures, as shown in Supplemental Figure 4A. Next, we observed a decrease in RIN4 protein stability over time in the presence of *Pst*-AvrRpt2 at both permissive and elevated temperatures in WT Col-0, as well as in the *ndr1* and *ics1-2* mutants (Figure 5A). Interestingly, in the *ndr1*/*35S*::*NDR1* over-expression line, we did not observe a reduction in RIN4 following infection with *Pst*-AvrRpt2, suggesting that overexpression of *NDR1* may protect RIN4 from cleavage. As expected, we observed no RIN4 disappearance following *Pst* inoculation over the same time frame (Supplemental Figure 4B).

**Figure 5.**
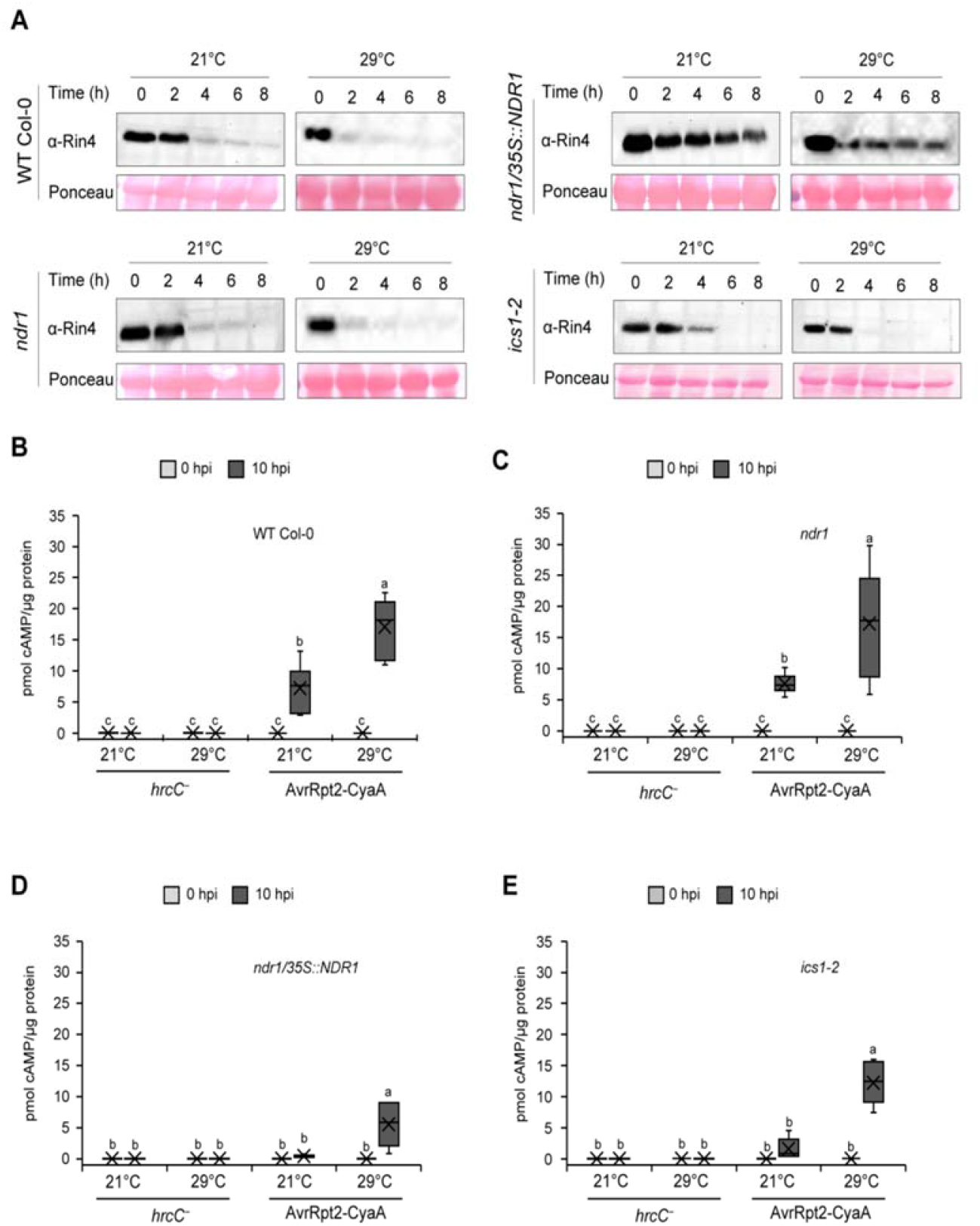
Overexpression of *NDR1* results in enhanced RIN4 stability in the presence of *Pst*-AvrRpt2. **(A)** Detection of RIN4 at 0, 2, 4, 6, and 8 hours after syringe-infiltration with *Pst*-AvrRpt2 (OD_600nm_ = 0.1) in wild-type (Col-0) and mutant plants. The total protein extracts were subjected to α -RIN4 western blot. Equal loading of protein was verified by ponceau S staining of the membrane after protein transfer. **(B), (C), (D)** and **(E)** Effector translocation in *ndr1, ndr1/35S::NDR1*, and *ics1-2*, respectively, following syringe-infiltrated with *Pst hrcC−* or *Pst* expressing AvrRpt2-CyaA (OD_600_=0.005). Tissue was collected at 0 and 10 hpi for quantification of cAMP which was normalized by total protein. Higher levels of cAMP indicate more translocation of bacterial effectors. *n* represents the total number of leaves from three independent biological repeats (n = 6). Values are plotted as box plots split by the median. Different letters represent a significant difference at *P* < 0.05 with Tukey’s honest significant difference (HSD) test. All data are representative of three independent experiments.

To gain a comprehensive understanding of the role of *NDR1* in ETI at elevated temperature, we further conducted disease assays in WT Col-0, *ndr1, ndr1*/*35S*::*NDR1*, and *ics1-2* plants following infection with *Pst* expressing, individually, the T3Es AvrRpm1 or AvrPphB. As shown, we observed comparable disease resistance to *Pst*-AvrRpt2 following both T3E infections in the *NDR1*-overexpression line at elevated temperature (Supplemental Figure 5 and 6). Based on these data, we surmise that enhanced resistance in *ndr1*/*35S*::*NDR1* against T3Es may be the result of their respective R protein interactions – both genetic and potentially physical – with NDR1, as well as via a yet defined function for NDR1 in SA-dependent signaling cascades.

To determine how overexpression of *NDR1* and the associated increase in SA might function in the activation of resistance, we next monitored the release of the negative regulation of immunity via the cleavage of RIN4 by the T3E cysteine protease AvrRpt2 (Axtell et al., 2003). To do this, we first evaluated the translocation of AvrRpt2 into plant cells via the type-III secretion system (T3SS) at 0, 6, and 10 h via the infection of plants with *Pst* expressing AvrRpt2 fused to a CyaA reporter (i.e., AvrRpt2-CyaA). As a negative control for these experiments, we employed a T3SS mutant *hrcC*^*−*^ carrying AvrRpt2-CyaA to ensure that detected *in planta* levels of CyaA arose via the action of a functional T3SS (Li et al., 2017). Additionally, *in planta* bacterial levels at each timepoint were quantified to eliminate any population-dependent translocation rate errors (Supplemental Figure 7). Consistent with previous studies that evaluated the impact of elevated temperatures on T3E translocation (Huot et al., 2017), we observed increased levels of cAMP at elevated temperature in all four plant genotypes compared to plants grown at permissive temperatures (i.e., 21°C) (Figure 5B-E). However, at 10 h, we observed the lowest levels of cAMP at elevated temperature in the *NDR1*-overexpression line compared to WT Col-0, followed by *ics1-2* and *ndr1* mutants (Figure 5B-E). This data is consistent with the increased stability of RIN4 in the *NDR1*-overexpression line (Figure 5A) and support a role for NDR1 in protecting RIN4 in the presence of *Pst-*AvrRpt2.

Previous work demonstrated that the plant defense inducer benzothiadiazone (BTH) induces pathogen resistance in a SA-dependent manner (Huot et al., 2017; Kouzai et al., 2018). To further uncouple the role of NDR1 and SA as a function of pathogen T3E activity, we first evaluated the effect of BTH on AvrRpt2-CyaA effector translocation in plants grown at both permissive and elevated temperatures. At neither temperature did we observe a significant change in the level of cAMP following BTH treatment, with the exception for in the *ics1-2* line; we hypothesize this that this is due to the low cAMP amount detected with mock treatment (Supplemental Figure 8A-B), which is likely due to subtle differences in buffer content (e.g., DMSO). Indeed, the lower levels of cAMP observed in BTH-treated *ics1-2* plants is consistent with our observations presented in Figure 5, wherein the amount of effector translocation was reduced in the *NDR1*-overexpression line, which also has increased levels of SA. As a control for these assays, we also enumerated *in planta* bacterial levels at 0 hpi to eliminate any population-dependent translocation rate errors (Supplemental Figure 9A-B). Overall, this result suggests that lower translocation rates observed in the *NDR1*-overexpression line (Figure 5D) could be a consequence of the elevated levels of SA in this line.

Having demonstrated the impact of SA on bacterial T3E translocation into the host cell, we next queried the role of SA on RIN4 cleavage by *Pst*-AvrRpt2, a function required for the robust activation of R-protein (e.g., RPS2) mediated ETI (Axtell and Staskawicz, 2003) following T3E (i.e., AvrRpt2) delivery and recognition. At the onset of this line of investigation, our working hypothesis was that given the enhanced level of resistance in the *NDR1*-overexpression line (Coppinger et al., 2004), cleavage of RIN4 by AvrRpt2 occurs more rapidly in this line, thereby leading to the release of negative regulation on the R-proteins RPS2 (Axtell and Staskawicz, 2003) and RPM1 (Mackey et al., 2003) and the robust activation of ETI. To test this, we first monitored the levels of RIN4 protein in BTH-treated plant lines hand-infiltrated with *Pst*-AvrRpt2. As shown in Supplemental Figure 8C, exogenous application of BTH did not protect RIN4 from cleavage by AvrRpt2 in WT Col-0, nor in the *ndr1* or *ics1-2* mutants. However, similar to results observed in Figure 5A, overexpression of *NDR1* did result in enhanced protection of RIN4 from cleavage by *Pst*-AvrRpt2. Based on this result, we conclude BTH induced SA does not protect RIN4 under *Pst*-AvrRpt2 treatment. Thus, the inability of SA to protect RIN4 from cleavage, coupled with the observed RIN4 protection in the *NDR1*-overexpression line, is likely due to the physical interaction, and stoichiometry of this association, between RIN4 and NDR1.

## DISCUSSION

Plant immune signaling during heat stress response has been described since the early 1900’s, wherein it was demonstrated that the spread of tobacco mosaic virus (TMV) necrotic lesions in *Nicotiana glutinosa*-infected leaves was more prevalent at elevated temperatures (Samuel, 1931). More than 75 years after this discovery, similar correlations have been described as they relate to the impact of elevated temperature on plant growth (Penfield, 2008), reproduction (McClung and Davis, 2010), and hormone signaling (Sakata et al., 2010). At a fundamental level, work using model plant-pathogen systems has demonstrated that plant resistance to bacterial pathogens is reduced under conditions of elevated temperature, a phenomenon that is hypothesized to be associated with the downregulation of SA-mediated immune signaling (Li et al., 2010; Huot et al., 2017). Collectively, these studies have provided foundational support for the ‘growth-defense’ paradigm - a model describing the allocation of cellular resources and the crosstalk between seemingly disparate genetic processes (Guo et al., 2018). In the current study, we sought to expand our understanding of the mechanisms that function at the nexus of heat stress response and immune signaling activation. To do this, we focused on the activation of a well-defined and genetically tractable immune signaling cascade, ETI. As presented herein, the data described point to the intersection of at least three immune signaling events (I. Transcriptional activation of immunity, II. Hormone response and signaling, and III. R Protein mediated immunity), each of which not only requires the function of NDR1 but is also impacted by the *Pst* type-III virulence effector AvrRpt2.

### Transcriptional activation of immunity: balancing abiotic and biotic stress

Previous studies have shown that plant response(s) to both biotic and abiotic stimuli are typically initiated by rapid, highly specific, changes in the transcriptional landscape; notably, the induction of genes associated with plant defense (Hu et al., 2012), and the attenuation of those required for growth and reproduction (Lee et al., 2014; Quint et al., 2016). Not surprisingly, these observations, as well numerous others, have led to the development of models that describe an important role for the co-regulation of processes that function antagonistically during simultaneous exposure to abiotic and biotic stressors (Hossain et al., 2018; Kim et al., 2020; Lu et al., 2021). In the current study, we sought to further the investigation of a role for rapid transcriptional changes in response to elevated temperature, through the lens of broader immune signaling responses. To do this, we generated 69 transcriptomes from four plant lines under permissive and elevated temperature, each of which have reported varied responses related to pathogen infection and hormone signaling (Century et al., 1997; Tao et al., 2003; Strawn et al., 2007; Catinot et al., 2008; Li et al., 2021). Through this approach, we identified two main clusters of DEGs that segregated based on genotype-independent expression, as well as those that were regulated in a genotype-specific manner under elevated temperature. Out of the two, the latter cluster was comprised of highly expressed genes in the *NDR1*-overexpression line. These genes were primarily related to GO terms like defense response and immune system function. Interestingly, we observed persistent stability in *RPS2* mRNA at elevated temperatures in the *NDR1*-overexpression line. Based on this, we hypothesized that in the absence of pathogen infection, the overexpression of *NDR1* leads to a preemptive transcriptional response to enhance resistance at elevated temperature. If true, the impetus for such a response would be the priming of immune signaling to protect plants during biotic stress in combination with abiotic stress exposure.

### Hormone response and signaling during abiotic and biotic stress perception

Phytohormones are an indispensable component of the plant immune system, required for the robust activation of both PTI and ETI (Miller et al., 2017; Yuan et al., 2021). For example, early work in this area has demonstrated that the immune signaling action of SA, which is synthesized via the isochorismate pathway, is rapidly produced following pathogen infection and required for downstream activation of immunity (Vlot et al., 2009). Interestingly, SA production is also affected when plants are exposed to both low (Li et al., 2020) and elevated (Huot et al., 2017) temperatures. As immune signaling modulators, previous work showed that RPS2 is also required not only for the production of SA, but also for the generation of pathogen-induced jasmonic acid (JA) and abscisic acid (ABA) production, supporting the hypothesis that to some degree, defense hormone production is ETI dependent (Liu et al., 2016).

In contrary to published data showing loss of virulent *Pst-*induced SA biosynthesis at elevated temperature, we observed that avirulent *Pst*-AvrRpt2 promotes SA synthesis in a temperature independent manner in WT Col-0 and the *NDR1-*overexpression line at 29°C (Huot et al., 2017). Intriguingly, SA levels were compromised in the *ndr1* mutant – similar to that detected in the SA deficient *ics1-2* mutant line at elevated temperature. Taken together, the stable expression level of *RPS2* upon heat stress prior to pathogen introduction, the high SA levels upon *Pst*-AvrRpt2 treatment with the inherently high SA and *NDR1* levels in the *NDR1*-overexpression line may have contributed to lowering the effector translocation enabling the RPS2 mediated ETI response in the absence of RIN4 depletion. The observed susceptibility in WT Col-0 despite the high levels of SA upon *Pst*-AvrRpt2 may be due to lower mRNA accumulation of *RPS2* compared to the NDR1 overexpression line at elevated temperature.

### Immune system stability

Recent studies aimed at identifying the underlying molecular-genetic mechanisms controlling immune signaling stability at elevated temperature have uncovered a relationship between an increase in temperature and the sustainable activity of host R-proteins (Venkatesh and Kang, 2019). For example, *Arabidopsis* plants subjected to a long-term (ca. 10 days) temperature acclimation at 28°C resulted in an approximate 8-fold increase in the *in planta* growth at 3 DAI of the virulent pathogen *Pst* compared to the plants grown at 22°C after 3 DAI. Furthermore, the same study also demonstrated that *Pst* expressing the T3Es AvrRpt2 and AvrRpm1 showed ten times more bacterial growth at 28°C compared to 22°C, indicating the plant defense responses mediated by *R*-genes are likely suppressed at higher temperatures (Wang et al., 2009), and/or are affected by the virulence activity of these effectors. In support of the former mechanism, it has been demonstrated that R-protein stability is linked to the presence of SA, which as noted above, plays an indispensable role in the plant defense response to bacterial pathogens. For example, the *R* gene-like Toll/interleukin-1 receptor (TIR)-NB-LRR type gene SUPPRESSOR OF *NPR1-1* CONSTITIUTIVE1 (*SNC1*) has emerged as a case-study for SA-dependent resistance signaling and a model defining the crosstalk between the *R* genes and hormones (Zhang et al., 2003; Yang and Hua, 2004). Interestingly, SNC1 protein accumulation is reduced at elevated temperatures, a phenomenon that is coincident with the reduction of SA at elevated temperatures (Zhu et al., 2010; Huot et al., 2017). Through a mutagenesis-based approach, coupled with the gain of function mutant *snc1-1* (Zhu et al., 2010) identified a temperature-insensitive mutant, *102snc1-1*, which retains pathogen resistance at both basal and elevated temperatures. This temperature insensitive immune response in the *102snc1-1* was further attributed to the high expression level of *PR1*, further supporting the involvement of SA (Zhu et al., 2010). Similarly, we observed sustained *PR1* mRNA accumulation/gene expression under elevated temperature in the *NDR1*-overexpression line. Indeed, current data supports a general model wherein R-protein-mediated resistance is suppressed at elevated temperatures together with an inhibition of SA (Huot et al., 2017). However, the precise mechanism(s) underpinning this suppression have remained elusive (Alcazar and Parker, 2011).

Growing evidence suggests that both nuclear localized TIR-NLRs and plasma membrane (PM) localized CC-NLR receptor mediated signaling pathways in a temperature-sensitive manner (Mang et al., 2012; Cheng et al., 2013). Unexpectedly, we found that in contrast to the reported suppression of RPS2 mediated ETI signaling at elevated temperatures, the *NDR1*-overexpression line remained resistant at elevated temperature. Previous work demonstrated that temperature acclimation at 32°C for 6 hours (i.e., short-term) did not impact the mRNA accumulation of key NB-LRR signaling components (e.g., *RPM1, RPS2, RIN4*, and *NDR1*) in the absence of pathogen infection (Cheng et al., 2013). Here, our data revealed the stable expression level of *RPS2* gene in the *NDR1*-overexpression line in the absence of pathogen under long-term (24 h) heat stress. This suggests that the early establishment of gene expression in key resistance components could be a preemptive strategy to defend against pathogen infection under conditions of environmental stress.

Cleavage of RIN4 by AvrRpt2 is required for the activation of RPS2-based ETI (Mackey et al., 2003). Previous results showed that RIN4 cleavage by AvrRpt2 occurs within ∼8 hpi at both 23°C and 32°C (Cheng et al., 2013). Consistent with this, we showed RIN4 degradation in WT Col-0, *ndr1*, and *ics1-2* at basal and elevated temperatures. However, in the *NDR1*-overexpression line, RIN4 remained protected from cleavage by AvrRpt2, suggesting a role of RIN4 protection conferred by the overexpression of *NDR1*. As previously reported, NDR1 and RIN4 physically interact *in planta* (Day et al., 2006; Knepper et al., 2011). Herein, we demonstrate that BTH-induced SA production was not sufficient to protect RIN4 from cleavage in WT Col-0, *ndr1*, and *ics1-2*, thereby eliminating the possibility of the involvement of the high levels of SA in the *NDR1*-overexpression line in the protection of RIN4. The simplest explanation for this is that overexpression of *NDR1*, and the interaction of NDR1 with RIN4, protects RIN4 from cleavage. What remains unclear is how *NDR1*-overexpression results in reduced effector translocation, an observation previously observed in *ics1-2* at both basal and elevated temperatures (van Dijk et al., 1999; Huot et al., 2017). As for the low effector translocation in the *NDR1*-overexpression line, two possible explanations exist. First, a loss of SA has been demonstrated to result in increased bacterial T3E translocation due to increased permeability of the cell in the *ics1-2* (Huot et al., 2017). Second, NDR1 has been shown to be required for PM-cell wall adhesion, and in the absence of *NDR1*, cells undergo rapid CaCl_2_-induced plasmolysis (Knepper et al., 2011).

One confounding observation presented herein is the increased stability – and/or protection – of RIN4 following *Pst*-AvrRpt2 treatment. This is surprising, given that overexpression of *NDR1* leads to enhanced resistance to *Pst*-AvRpt2 (Coppinger et al., 2004). To explain this, we posit that only a minor amount of RIN4 cleavage is required for full activation of ETI, a mechanism that ensures the robust activation of resistance following release of RIN4 negative regulation. An alternative, or concomitant, explanation is that high levels of SA, together with the stable expression of *RPS2*, are responsible for the observed resistance in the *NDR1*-overexpression line. In this vein, it is interesting to hypothesize that one function of NDR1 is that of a cellular homeostasis sensor, functioning as a component of a general cellular surveillance (Battaglia et al., 2008; Knepper et al., 2011).

Related to the data presented herein, the observed temperature sensitivity of *NDR1, RPS2*, and other defense genes, and the increase of bacterial effector translocation in WT Col-0 and mutant plants at elevated temperature supports the disease progression upon pathogen entry. We propose that the rescue of ETI in the absence of RIN4 degradation in the *NDR1*-overexpression line is potentially due to a complex interaction involved the stability of *RPS2* mRNA expression and a decrease in T3E translocation rates at elevated temperature (Figure 6). While much work remains towards fully defining the role of NDR1 at the nexus of biotic and abiotic signaling, the data described provides insight into a role for NDR1 as a stabilizing component, and potential scaffolding mechanism, required for the maintenance of homeostasis during abiotic and biotic stress response signaling.

**Figure 6.**
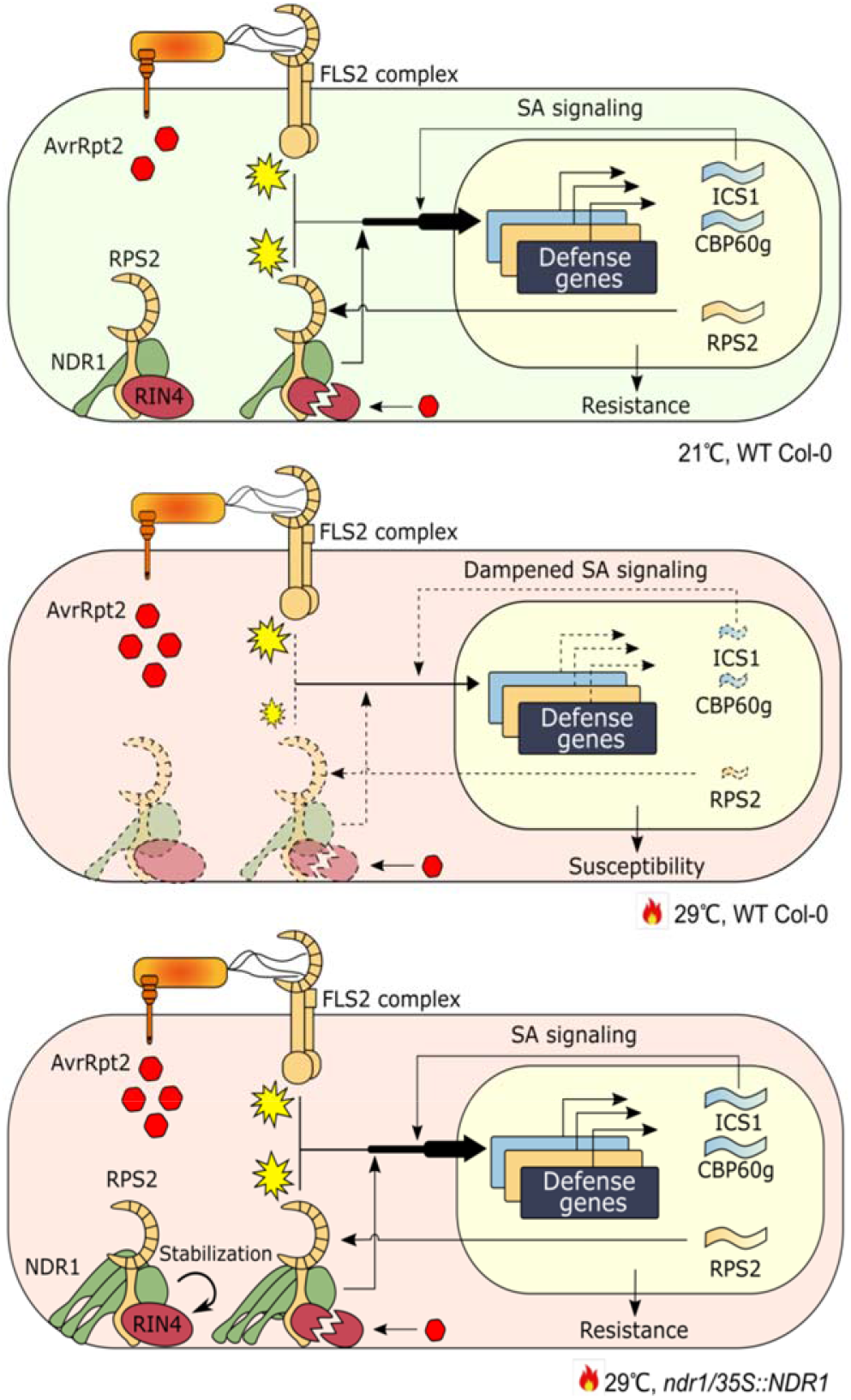
The schematic diagram of the mechanism of rescued ETI in elevated temperature by overexpressed *NDR1*. Dashed lines indicate dampened signaling pathway. In plant immunity, NDR1 genetically interacts with RPS2 and RIN4 to facilitate ETI in response to AvrRpt2. Concomitantly, NDR1 contributes to a robust pro-immune transcriptome, including the upregulation of *RPS2* and genes involved in SA signaling. At elevated temperature (29°C), transcription of these defense genes is inhibited, rendering plants susceptible to bacteria pathogen infection. In the *ndr1/35S::NDR1* overexpression line, increased levels of *NDR1* rescues the transcription of *ICS1* and *CBP60g* under elevated temperatures, thus sustaining the production of SA and its signaling pathway. In parallel, *NDR1*-overexpression enhances *RPS2* mRNA accumulation and stabilizes RIN4, the guardee, to sustain the function of the NB-LRR complex.

## METHODS

### Plant Material and Growth Conditions

*Arabidopsis thaliana* wild-type (WT) Columbia-0 (Col-0) plants, and mutant plants (*ndr1, ndr1/35S::NDR1*, and *ics1-2*), all of which are in the Col-0 background, were used in this study. Plants were grown in Arabidopsis soil mix comprised of equal parts of Sure-Mix (Sure, Galesburg, Michigan), Perlite (PVP Industries, Ohio), and Vermiculite (PVP Industries). Plants were grown for 3-4 weeks at 21°C under a 12 h/12 h light/dark cycle with 60% relative humidity and a light intensity of 120 μmol photons m^-2^s^-1^ prior to heat stress in a BigFoot Series growth chamber (BioChambers). For temperature assays, plants were separated into two chambers set to either 21°C (permissive) or 29°C (elevated). Plants were subjected to heat stress treatment for 48 h at 29°C prior to pathogen treatment. The area of infiltration was marked to ensure that the leaf tissue subsequently collected for the assays contained bacterial inoculum.

### Bacterial Strains and Disease Assays

*Pseudomonas syringae* pv. *tomato* (*Pst*) DC3000 harboring the open-reading frames of the T3E; AvrRpt2, AvrPphB, AvrRpm1, as well as the empty vector (EV; pVSP61; (Kunkel et al., 1993)) were grown on NYGA (5 g/L Bacto-peptone, 3 g L^-1^ yeast extract, and 20 mL L^-1^ glycerol, with 15 g L^-1^ agar for solid medium) containing 25 μg mL^-1^ kanamycin (kan) and 100 μg/mL rifampicin (rif) for 2 days at 28°C. After 48 h, bacterial cultures were resuspended in 5 mM MgCl_2_ at the desired concentrations for *in planta* growth assays. The same growing and culture suspension method was followed for the CyaA fusion protein tagged variant of AvrRpt2 and T3SS mutant *hrcC*^*−*^, with the exception that NYGA plates contained 25 μg mL^-1^ rif and 10 μg mL^-1^ gentamycin (gen).

### *In planta* Bacterial Growth Assays

*Pst* harboring AvrRpt2, AvrPphB, AvrRpm1, and EV (empty vector; control) inoculums were prepared at an optical density (OD) at 600nm = 0.0005 (ca. 5×10^5^ CFU mL^-1^), OD_600nm_ = 0.1 (ca. 10^8^ CFU mL^-1^) and OD_600nm_ = 0.0075 (ca. 7.5×10^6^ CFU mL^-1^) for growth curve, HR and BTH treatment assays, respectively. Bacterial inoculations were performed on multiple (n > 3) fully expanded leaves from 3-week-old Arabidopsis plants grown at permissive (21°C) and elevated (29°C) temperatures. Plants were inoculated with *Pst* isolates using a needleless syringe. Three biological replicates were performed for each assay. For *in planta* bacterial growth curve analyses, 3 mm leaf disks from 3 plants (3 leaves per sample) were collected at 3 DAI. Harvested leaf discs were incubated in 5 mM MgCl_2_ + 0.1% Tween-20 at 28°C, on a rotary platform shaker, for 1 h. After 1 h, each sample was serially diluted (10-fold increments) and 5 μL of each dilution was plated on NGYA plates containing ½-strength antibiotics (i.e., rif and kan). After 2 d incubation at 28°C, bacterial CFUs were counted. For HR analysis, infected leaves (24 hpi) were collected, and phenotypes were recorded by digital photography.

### RIN4 Western Blot Analysis

Two leaves from each plant genotype were hand infiltrated (OD_600nm_ = 0.1; ca. 10^8^ CFU/mL) with *Pst* expressing either AvrRpt2 or EV. Infiltrated leaves were collected at the designated timepoints, placed into a sterile 2 mL centrifuge tube, and were flash frozen in liquid nitrogen. Samples were ground in extraction buffer (20 mM Tris pH 7.5, 150 mM NaCl, 1 mM EDTA, % Triton X-100, 0.1% SDS, 5 mM DTT, 10X Sigma protease inhibitor mixture) and centrifuged at 20,000 x *g* for 10 min at 4°C. After centrifugation, the supernatant was collected as the total protein extract. Total protein of 50 μg was equalized using 6X loading buffer (0.375M Tris pH 6.8, 12% SDS, 60% glycerol, 0.6M DTT, 0.06% bromophenol blue) as a dilutant. Samples were separated by SDS-PAGE using 4-12% NuPAGE gels (Invitrogen) followed by transfer onto nitrocellulose membranes (GVS North America) for western blot analysis.

To generate antibodies to RIN4, the open reading frame of *RIN4* was cloned into pENTR and then moved into the destination protein expression Gateway pDEST17 vector (ThermoFisher, Cat# 11803012) using LR Clonase enzyme mix (ThermoFisher, Cat# 11791043). RIN4 was expressed in *E. coli* C41 (DE3) cells (Lucigen, Cat# 60442-1) and purified using Ni-NTA agarose (Qiagen, Cat# 30210) according to the vendor’s instructions. Purified total protein extracts were separated using NuPAGE 4-12% Bis-Tris gels and the band containing the majority of RIN4 protein was excised for antibody production. Polyclonal rabbit anti-RIN4 antibody was produced by Cocalico Biologicals, Inc. The specificity of anti-RIN4 antibody was confirmed by western blot analysis using WT Col-0 and *rps2/rin4* mutant plant lines, as well as transient expression of RIN4 protein in *N. benthamiana*. Anti-RIN4 sera was used at a concentration of 1:5000 in 1X TBST (1 M Tris pH 8.0, 1% Tween 20, 5% dehydrated milk).

### Phytohormone Analysis

Leaves were infiltrated with *Pst* suspended in 5 mM MgCl_2_ expressing AvrRpt2, AvrRpm1, or *Pst* harboring pVSP61 (EV) using a 1 mL needleless syringe at a concentration of OD_600nm_ = 0.0005 (ca. 5×10^5^ CFU mL^-1^). Mock inoculation controls were performed using 5 mM MgCl_2_. Quantification of phytohormones was performed as previously described (Velasquez et al., 2017), with minor modifications. In brief, fifty mg fresh weight (2-3 leaves, 3-to-4-week-old plants) were transferred to 2 mL sterile centrifuge tubes, flash frozen in liquid nitrogen, and stored at -80°C until processing.

For hormone extraction and quantitative evaluation, frozen tissue was ground using a TissueLyser II (Qiagen) and incubated on a rocking platform at 4°C for 24 h in extraction buffer (80:20 v/v HPLC-grade methanol:water with 0.1% formic acid (v/v), 0.1 g L^-1^ butylated hydroxytoluene). Samples were centrifuged at 12,000 x *g* for 10 min at 4°C and resultant supernatants were collected and were filtered through a 0.2 mm PTFE (polytetrafluoroethylene) membrane (Millipore).

Abscisic acid (ABA)-d_6_ (Toronto Research Chemicals Inc,) served as an internal standard. Injections of plant extracts (10 mL per injection) were separated on a Waters Acquity BEH-C18 column (2.1 × 50 mm, 1.7 mm) installed in the column heater of an Acquity Ultra Performance Liquid Chromatography (UPLC) system (Waters Corporation). A gradient of 0.1% aqueous formic acid (solvent A) and methanol (solvent B) was applied in a 5-min program with a mobile phase flow rate of 0.4 mL min^-1^ as follows: 0 to 0.5 min hold at 98% A and 2% B, transition to 70% B at 3 min, to 99% B at 4 min, hold at 99% B to 5 min, return to 98% A at 5.01 min and hold at 98% A to 6 min. The column was maintained at 40°C and interfaced to a Waters Xevo TQ-XS mass spectrometer equipped with electrospray ionization and operated in negative ion mode with a capillary voltage of 1.00 kV. The flow rates of cone gas and desolvation gas were 150 and 800 L h^-1^, respectively. The source temperature was 150°C and the desolvation temperature was 400°C. Collision energies and source cone potentials were optimized for each compound using QuanOptimize software (Waters Corporation). Peak areas were integrated, and the analytes were quantified based on standard curves generated from peak area ratios of analytes. Data acquisition and processing was performed using Masslynx 4.1 software (Waters Corporation). Analytes were quantified by converting peak area to phytohormone concentration (nM) per gram of dry weight of leaf tissue using a standard curve specific to each compound.

### Adenylate Cyclase (CyaA) Assay

To monitor *Pst* type-III effector delivery, a CyaA assay was performed as previously described (Fu et al., 2006; Chakravarthy et al., 2017), with slight modification. In brief, leaves from 4-week-old Arabidopsis plants were infiltrated with *Pst* expressing AvrRpt2-CyaA or the T3SS mutant *hrcC*^*−*^carrying AvrRpt2-CyaA suspended in 5 mM MgCl_2_ at a concentration of OD_600nm_ = 0.005 (ca. 5 × 10^6^ CFU cm^-1^) using a 1-mL needleless syringe. Leaf samples were harvested from 2 plants (2 leaves per sample) at 0, 6, and 10 hpi and snap frozen in liquid nitrogen. Cyclic adenosine monophosphate (cAMP) levels were quantified using the direct cAMP ELISA kit (Enzo Life Sciences, ADI-900-066).

### RNA Extraction, Library Preparation, and RNA Sequencing

RNA-seq analyses were performed on Arabidopsis plants representing four genotypes: WT Col-0, *ndr1, ndr1/35S::NDR1*, and *ics1-1*. Plants were grown at permissive temperatures (i.e., 21°C) for 23 days and then moved to elevated (i.e., 29°C) temperatures. Upon moving to 29°C, two fully expanded leaves from four different plants (8 leaves) were harvested as a single biological replicate at 0, 6, and 24 h. Tissue isolations were collected from three independent experimental replications, each containing three biological replicates. Total RNA was extracted using the RNeasy Plant Mini kit (Qiagen). DNA was removed from the sample by using TURBO DNA-Free™ kit (Thermo-Fisher). RNA samples were quantified using Nanodrop 2000 spectrophotometer (Thermo-Fisher).

### Construction of Strand-Specific RNA-seq Libraries

Construction of the RNA-seq libraries and sequencing on the Illumina NovaSeq 6000 were performed at the Roy J. Carver Biotechnology Center at the University of Illinois at Urbana-Champaign. Total RNAs were run on a Fragment Analyzer (Agilent, CA) to evaluate RNA integrity. RNA-seq libraries were constructed with the TruSeq Stranded messenger RNAs (mRNA) Sample Prep kit (Illumina). Briefly, polyadenylated mRNAs were enriched from 500 ng of high-quality DNA-free total RNA with oligo-dT beads. The mRNAs were chemically fragmented, annealed with a random hexamer, and converted to double stranded cDNAs, which were subsequently blunt-ended, 3′-end A-tailed, and ligated to indexed adaptors. Each library was ligated to a uniquely dual indexed adaptor (unique dual indexes) to prevent index switching. The adaptor-ligated double-stranded cDNAs were amplified by PCR for 8 cycles with the Kapa HiFi polymerase (Roche, CA) to reduce the likeliness of multiple identical reads due to preferential amplification. The final libraries were quantitated with Qubit (Thermo-Fisher) and the average library fragment length was determined on a Fragment Analyzer. The libraries were diluted to 10nM and further quantitated by qPCR on a CFX Connect Real-Time qPCR system (BioRad) for accurate pooling of the barcoded libraries and maximization of number of clusters in the flow cell. A total of 90 RNA-seq libraries were prepared from 1 μg of total RNA.

### Sequencing of Libraries on the NovaSeq Instrument

The pooled barcoded RNA-Seq libraries were loaded on a NovaSeq S2 lane for cluster formation and sequencing. The libraries were sequenced from one end of the fragments for a total of 100nt. The fastq read files were generated and demultiplexed with the bcl2fastq v2.20 Conversion Software (Illumina).

### Expression and Differential Analysis

The adapter sequences and low-quality bases (q < 10) were trimmed by Trimmomatic (Bolger et al., 2014). Resultant cleaned reads were mapped to the TAIR10 reference genome using HISAT2 (Kim et al., 2015). Mapped read counts for each gene were generated using the Htseq (Anders et al., 2015) command. The statistical analysis of the RNA-seq data was performed in the R environment (version 4.0.5). Mitochondrial and chloroplast genes were excluded, as they are not poly (A)-tailed. Genes with mean read counts of fewer than 10 per library were considered to be expressed at low levels and were excluded from the analysis. The resulting count data were subjected to trimmed-mean of M-values normalization using the function calcNormFactors in the package edgeR, followed by log-transformation by the function voomWithQualityWeights in the package limma to yield log_2_ counts per million. To each gene, a linear model was applied using the lmFit function in the limma package with the following terms: Sgetr = GETget+ εgetr, where S, log_2_ expression value, GET (genotype:environment:time) interaction, r, biological replicate, and ε, residual. For variance shrinkage in the calculation of p-values, the eBayes function in the limma package was used. Next, the resulting p-values were corrected for multiple hypothesis testing by calculating the Storey’s q-values using the function qvalue in the package qvalue. To extract genes with significant expression changes, the cutoff of q-value < 0.01 and > 2 fold expression changes were applied. AgriGO was used for GO enrichment analysis with default settings (Du et al., 2010). To create heatmaps, average linkage hierarchical clustering with uncentered Pearson correlation as a distance measure was carried out using CLUSTER 3.0 (Eisen et al., 1998) followed by visualization using TREEVIEW (Eisen et al., 1998).

Coexpression network analysis was performed using the R package WGCNA (Langfelder and Horvath, 2008). Genes with little expression variances (< 0.2) across the samples were excluded. Normalized and log_2_-transformed read counts of the resulting 11898 genes were used for constructing a singed hybrid network. The adjacency matrix was constructed using the adjacency function with the power of 14, and the topological overlap was then calculated from the adjacency matrix using the TOMsimilarity function. Average linkage hierarchical clustering was applied to the topological overlap for grouping genes with highly similar coexpression relationships. The Dynamic Hybrid Tree Cut algorithm was used to cut the hierarchal clustering tree, and 37 modules were defined as branches from the tree cutting. For the construction of the NDR1-centered network, eigengene-based gene connectivity, kME, was calculated using the signedKME function to select co-expression modules whose expression patterns are highly correlated to that of NDR1 (|kME| > 0.6). The relationships of these modules with *NDR1* were visualized using Cytoscape (Shannon et al., 2003).

### Accession Numbers

The RNA-seq data generated herein are contained within the National Center for Biotechnology Information (NCBI) Short Read Archive (SRA). The Illumina RNA-seq reads were deposited in BioProject under project ID: PRJNA778239.

## Supplemental Data

**Supplemental Figure 1**. Temporal dynamics of transcriptome responses to heat stress.

**Supplemental Figure 2**. Expression patterns of coexpression modules.

**Supplemental Figure 3**. Disease assays at 0 hpi after avirulent phytopathogen bacteria treatment.

**Supplemental Figure 4**. Temporal detection of RIN4 after virulent *Pst* treatment.

**Supplemental Figure 5**. Hypersensitive response and disease phenotypes after treatment with *Pst*-AvrRpm1 and *Pst*-AvrPphB.

**Supplemental Figure 6**. Disease assays at 3 dpi after avirulent phytopathogen bacteria treatment.

**Supplemental Figure 7**. Bacterial growth after syringe-infiltration with *Pst* expressing *hrcC™* and AvRpt2-CyaA in wild-type (Col-0) and mutant plants.

**Supplemental Figure 8**. Benzothiadiazole (BTH) induced SA does not protect RIN4 under *Pst*-AvrRpt2 treatment.

**Supplemental Figure 9**. Bacterial growth at 0 hpi in BTH untreated wild-type (Col-0) and mutant plants after syringe-infiltration with mock and *Pst* expressing AvrRpt2-Cya.

**Supplemental Data Set 1**. Mean centered expression levels (log_2_) of differentially expressed genes.

**Supplemental Data Set 2**. Differentially expressed genes in Cluster I.

**Supplemental Data Set 3**. Gene ontology enrichment of Cluster I containing genotype-independent temperature responsive genes (FDR < 0.05).

**Supplemental Data Set 4**. Differentially-expressed genes in Cluster II.

**Supplemental Data Set 5**. Gene ontology enrichment of Cluster II containing genotype specific temperature responsive genes (FDR < 0.05).

**Supplemental Data Set 6**. The number of upregulated genes between genotypes.

**Supplemental Data Set 7**. The number of downregulated genes between genotypes.

**Supplemental Data Set 8**. GO enrichment analysis of the genes in each co-expression module (FDR < 0.05).

**Supplemental Data Set 9**. Description of the genes in each co-expression module.

## ACKNOWLEDGEMENTS

We would like to acknowledge the support of the MSU Plant Resilience Institute for providing funding to the laboratory of B.D. Research in the laboratory of B.D. is supported by the National Institutes of General Medical Sciences (1R01GM125743) and the National Science Foundation-National Institute of Food and Agriculture (USDA) joint Plant-Biotic Interactions Program (IOS-1146128). Research in the laboratory of A.M. is supported by JST PRESTO (JPMJPR17Q6), Grant-in-Aid for Scientific Research (B) (19H02960), and Grant-in-Aid for Scientific Research on Innovative Areas (B) (21H05151). Research in the laboratory of K.T. is supported by the Fundamental Research Funds for the Central Universities (Program No. 2662020ZKPY009), the Huazhong Agricultural University Scientific & Technological Self-innovation Foundation, and Joint Funding of Huazhong Agricultural University and Agricultural Genomics Institute at Shenzhen, Chinese Academy of Agricultural Sciences (SZYJY2021007). We would also like to thank Masaki Shimono for experimental assistance, Sheng Yang He for providing the *ics1-2* seeds and Noel Day for editorial comments.

## AUTHOR CONTRIBUTIONS

Designed framework: S.P.S. and B.D. Conducted experiments: S.P.S., Y.-J.L., Y.K., Analyzed data: S.P.S., H.C., Y.-J.L., P.L., Y.K., A.M., K.T., and B.D. Wrote the manuscript: S.P.S. and B.D. All authors provided comments and editorial input during manuscript preparation and revision.

